# G3BP1 Maintains Vascular Endothelial Barrier Integrity by Negatively Regulating the MYD88-ARNO-ARF6 Signaling Pathway

**DOI:** 10.1101/2024.09.09.612151

**Authors:** Weiyue Sun, Haoran Wu, Yuxi He, Huiqiao Chen, Yuanhui Meng, Guofang Tang, Jinshun Zhu, Zhengwang Wen, Rongzhou Wu, Guowei Wu, Chunxiang Zhang, Maoping Chu, Bin Wen

## Abstract

Endothelial permeability is essential for vascular function. This process is regulated by intercellular junctions, including adherens junctions (AJs) and tight junctions (TJs), which form the endothelial barrier and dynamically connect adjacent cells. However, the mechanisms that control endothelial permeability are not fully understood.

We investigated the role of the RNA-binding protein G3BP1 in regulating endothelial permeability. We examined the effects of Loss of g3bp1 in conditional knockout mice and primary human HUVEC cells. We analyzed endothelial barrier integrity and permeability, focusing on AJ and TJ expression, under both normal and LPS-induced inflammatory conditions.

Loss of g3bp1 in vascular endothelial cells decreased AJ and TJ expression, compromised barrier integrity, and increased permeability. These effects were exacerbated under LPS-induced inflammatory conditions. Loss of g3bp1 also increased the expression of MYD88, ARNO, and ARF6, and enhanced ARF6 activity. Mechanistically, G3BP1 bound to and stabilized MYD88 mRNA, negatively regulating the MYD88-ARNO-ARF6 signaling pathway. Inhibiting this pathway by reducing MYD88 or ARF6 expression, or inhibiting ARNO activity, restored AJ/TJ expression and barrier function in G3BP1-deficient models.

Our findings identify G3BP1 as a novel regulator of vascular endothelial permeability. G3BP1 acts by negatively regulating the MYD88-ARNO-ARF6 signaling pathway. Targeting G3BP1 may be a potential therapeutic strategy for diseases associated with endothelial barrier dysfunction.

## 1. Introduction

Endothelial permeability is critical for normal vascular physiology and organ function[1]. Dysregulation of endothelial permeability causes or exacerbates a variety of human pathologies including inflammatory airway diseases, arthritis, chronic bowel disease, cancer, infections, and ischemic stroke, thereby influencing disease morbidity and treatment [2].

Endothelial permeability is governed by a barrier formed by intercellular junctions, including adherens junctions (AJs) and tight junctions (TJs) [2]. The assembly, stability, and function of AJ/TJ are regulated by various modulators and signaling pathways, such as PKC[3–5], RhoA/ROCK[6], MAPK[7–10], VEGF[11], PI3K/mTOR[12], and MYD88-ARNO-ARF6 signaling[13, 14]. These pathways influence the expression and organization of AJs and TJs through transcriptional and post-translational mechanisms involving transcription factors, kinases, phosphatases, and small GTPases. Despite considerable progress in understanding endothelial barrier regulation, our knowledge remains incomplete. The identification of novel molecular players in endothelial barrier regulation is crucial for developing targeted therapies for permeability-related disorders.

Recently, RNA-binding proteins (RBPs) have emerged as critical regulators of gene expression and cellular function in cardiovascular health and disease [13]; however, their role in the regulation of endothelial barrier remains largely unexplored. Over 300 RBPs have been identified in various cardiovascular cell types, including endothelial cells [15]. However, only a handful have been linked to endothelial barrier regulation. For instance, QKI affects VE-cadherin and beta-catenin protein expression and maintains endothelial barrier function [16]. These findings underscore the need for further investigation into the contributions of RBPs to endothelial barrier regulation.

G3BP1 is a heterogeneous nuclear multi-functional RBP. G3BP1 regulates gene expression and stress granule formation [17] by controlling mRNA stability[18–24] and translation[25, 26]. Additionally, G3BP1 participates in various signaling pathways through protein interactions [27–30]. G3BP1 dysfunction is implicated in various diseases, including cancer [31], immune responses [19, 27] , and neuronal regeneration [25].

Although G3BP1’s role in cancer, antiviral responses, and neuronal regeneration are well-established, its function in vascular physiology and pathology is less understood. Emerging evidence points to its potential involvement in the vascular system. For example, G3BP1 plays a crucial role in mediating Wnt signaling in aortic smooth muscle cells, influencing NFATc4 transcription and potentially contributing to atherosclerosis [29]. In vascular endothelial cells, G3BP1 regulates glucocorticoid receptor-mediated miRNA maturation [32]. Intriguingly, global *G3bp1* knockout mice exhibit intracranial hemorrhage during embryonic development [20], suggesting a potential role for G3BP1 in maintaining vascular integrity.

Despite this suggestive evidence, the precise role of G3BP1 in vascular integrity remains elusive. We hypothesized that G3BP1 maintains vascular endothelial barrier integrity by negatively regulating the MYD88-ARNO-ARF6 signaling pathway. To test this hypothesis, we examined the effects of G3BP1 deficiency on barrier integrity and the expression and activity of the MYD88-ARNO-ARF6 signaling pathway in lung vascular endothelial cells of endothelial-specific *G3bp1* knockout mice and *G3BP1*-depleted human umbilical vein endothelial cells (HUVECs).

## 2. Materials and Methods

### 2.1. Generation of G3BP1 conditional KO mice

*G3bp1*-floxed mice were generated using CRISPR/Cas9 technology at the Shanghai Model Organisms Center. The generation process involved microinjecting C57BL/6 fertilized eggs with a donor vector, Cas9 mRNA, and guide RNAs targeting G3bp1 introns 2 and 3. Founder mice were backcrossed with wild-type C57BL/6 mice to obtain heterozygous G3bp1 flox/+ mice. These were then self-crossed to produce homozygous *G3bp1 ^flox/flox^* mice. These mice were crossed with B6;129-Tg(Cdh5-cre) Smoc mice (Stock No. NMX-TG-192013) to generate *Cdh5:Cre*; *G3bp1 ^flox/flox^* mice. Genotyping was performed using PCR on tail genomic DNA with the following primer pair: forward (5’-ctgtccccactgccttcag-3’) and reverse (5’-gcttggtactcccgctc-3’). All animal protocols adhered to guidelines approved by the Ethics Committee of Wenzhou Medical University. Mice were housed in a specific pathogen-free (SPF) facility under standard conditions.

### 2.2. Generation of constructs

*G3BP1* was amplified by RT-PCR from primary HUVEC cDNA using the following primers:Forward: (5’- ATGGTGATGGAGAAGCCTAG -3’), and Reverse: (5’- GCAGTTTATAACAGACTGGG -3’). The amplified G3BP1 CDS was then ligated into the BamHI and EcoRI sites of the pCMV vector. All constructs were verified by sequencing.

### 2.3. Antibodies and reagents

The following antibodies were used for immunostaining: VE-cadherin (1:50, sc-9989; Santa cruz), p120 (1: 50, sc-23873;Santa cruz), CD31 (1: 100, AF3628; R&D system), G3BP1 (1:1000,13057-2-AP;Proteintech), Alexa Fluor 488 donkey anti-rabbit antibody (1:200, A-21206; invitrogen) or Alexa Fluor 594 donkey anti-goat antibody (1:200,A-21209; invitrogen). The following antibodies were used for western blotting: VE-cadherin (1: 1000, #2158; Cell Signaling Technology), p120 (1: 200,sc-23873;Santa cruz), ZO-1(1:1000, Cell Signaling Technology,5406), Claudin5 (1:200, #49564; Cell Signaling Technology), ARF6 (1:1000, 20225-1-AP; Proteintech), MYD88 (1:1000, #3699; Cell Signaling Technology), Cytohesin2 (1:1000, 10405-1-AP; Proteintech), β-actin ( 1:1000, #4967, Cell Signaling Technology), G3BP1 (1: 100, 66486-1-Ig; Cell Signaling Technology) . G3BP1 RIP was performed using an antibody against G3BP1 (1:1000, RN048PW; Medical Biological Laboratories).

G3BP1 and MyD88 siRNAs were purchased from GenePharma Technology (Shanghai, China). LPS (L2630; Sigma-Aldrich) was dissolved in sterile PBS for cell culture experiments and in saline for mouse experiments. NAV2729 (HY-112473; MedchemExpress) was dissolved in dimethyl sulfoxide (DMSO) to prepare a stock solution and used at a final concentration of 10μM. TJ-M2010-5 (TJM; HY-139397; MedchemExpress) was prepared in a solution of 5% DMSO, 40% PEG300, 5% Tween 80, and 50% ddH2O to a final concentration of 15 mg/kg.

### 2.4. Miles Assay

Mice were injected with PBS or lipopolysaccharide (LPS, 10 mg/kg, intraperitoneally) for 6 hours, followed by EBD (30 ml/kg, tail vein injection) for 1 hour. After euthanasia, multiple organs including lungs and small intestines were collected, imaged, weighed, and homogenized in phosphate-buffered saline (PBS). Homogenates were incubated with formamide at 60°C for 48 hours, centrifuged, and the optical density of the supernatant was measured at 610 nm. EBD concentration was calculated using a standard curve (micrograms EBD per gram tissue).

### 2.5. Wet weight/dry weight ratio of lungs

Mice were anesthetized with 4% chloral hydrate (0.1 ml/kg, intraperitoneal injection). The right lung was isolated and immediately weighed. The lung was then dried at 70° C for 72 hours, after which the dry weight was recorded. The wet weight/dry weight ratio was calculated to assess pulmonary edema.

### 2.6. Histological evaluation

Lung tissues from the *G3bp1 ^flox/flox^* and *Cdh5:Cre*; *G3bp1 ^flox/flox^* mice were fixed in 4% paraformaldehyde. The tissues were then embedded in paraffin, and sectioned at 5μ m thickness. Sections were stained with hematoxylin and eosin (H&E) and examined using a bright-field microscope (OLYMPUS, BX43) at 200 x and 400 x magnification. The evaluation focused on alveolar wall thickening and neutrophil infiltration in the alveolar and interstitial spaces. Inflammatory cell infiltration in the lung was scored using the following criteria:

- points: No inflammatory cell infiltration.
- point: Mild inflammatory cell infiltration.
- points: Increased inflammatory cell infiltration with uneven distribution.
- points: Extensive inflammatory cell infiltration, evenly distributed but not aggregated into clusters.
- points: Severe inflammatory cell infiltration with aggregation into clusters.

### 2.7. Transmission Electron Microscopy

Mice were anesthetized with ketamine and xylazine (250 and 20 mg/kg, respectively, intraperitoneally). Lungs were perfused and fixed with 2.5% glutaraldehyde in cacodylate buffer, post-fixed in 1% osmium tetroxide, dehydrated through an ethanol series, and embedded in epoxy resin. Ultrathin sections were prepared, stained with uranyl acetate and lead citrate, and examined under a electron microscope (Hitachi H-7700) at 80 kV.

### 2.8. Cell culture and transfection

HUVECs were purchased from ScienCell Research Laboratories (#8000; ScienCell ) and cultured in endothelial growth medium (#1001; ScienCell) supplemented with 5% fetal bovine serum (Lonza, Basel, Switzerland) at 37°C in a 5% CO2 atmosphere. Experiments were conducted using cells between passages 4 and 8.

For siRNA knockdown experiments, HUVECs were transfected with Scramble or G3BP1 siRNA using Lipofectamine™ RNAiMAX (ThermoFisher) according to the manufacturer’s instructions. Briefly, HUVECs were seeded in 12-well plates at a density of 4 × 10^4 cells/well. For siRNA transfection, 20 pmol of siRNA was diluted in 250 μL of Opti-MEM Reduced Serum Medium. Separately, 2.5 μL of Lipofectamine RNAiMAX was diluted in 250 μL of Opti-MEM. The diluted siRNA was then combined with the diluted Lipofectamine RNAiMAX, mixed gently, and incubated for 10-20 minutes at RT. This mixture was then added to each well containing 1 mL of culture medium. The sequences of G3BP1 siRNAs were: si*G3BP1*-1 (5’-CATTAACAGTGGTGGGAAA-3’) and si*G3BP1*-2 (5’-AGGCTTTGAGGAGATTCAT-3’).

For overexpression experiments, HUVECs were transfected with GFP expression vector (control) or G3BP1 expression vector using PolyJet reagent (SignaGen Laboratories). Briefly, 1μg of plasmid DNA was mixed with 3μL of PolyJet reagent and added to 60% confluent HUVECs in each well of a 6-well plate according to the manufacturer’s instructions. Cells were harvested 72 hours after transfection, and lysates were prepared by homogenization in 2X Laemmli buffer (Sigma-Aldrich).

### 2.9. In vitro FITC-dextran permeability assay

HUVECs were cultured in transwell chambers with 0.4 µm pore size polyester membranes (Corning, USA) until 70% confluence. After treatment with Scramble or G3BP1 siRNA for 12 hours, FITC-dextran (40 kDa) in serum-free medium was added to the upper chamber. After 8 hours, fluorescence in the lower chamber was measured with a microplate reader (Ex = 492 nm; Em = 520 nm).

## 3.0. Immunostaining

For cell immunostaining, HUVECs on glass coverslips were fixed with 4% paraformaldehyde for 15 minutes at RT and then permeabilized with 0.1% Triton X-100 for 10 minutes. Blocking was performed with 5% BSA in PBS for 1 hour. Cells were incubated overnight at 4°C with primary antibodies against VE-cadherin and ZO-1. After washing with PBS, cells were incubated with appropriate fluorescently labeled secondary antibodies for 1 hour at RT in the dark. Nuclei were counterstained with DAPI (1:1000) for 5 minutes. Coverslips were mounted onto glass slides using an anti-fade mounting medium.

For section immunostaining, paraffin embedded sections fixed with formalin were deparaffinized with xylene, and hydrate through a graded series of alcohols. Antigen retrieval was done by bringing the slides to a boil in 10 mM sodium citrate buffer (pH 6.0) and then maintain at a sub-boiling temperature for 10 minutes. Sections were washed twice for 10 minutes in PBS with 0.3% Triton X-100 to permeabilize the tissue. After blocking with 2% normal donkey serum in PBST for 1 hour at RT, sections were incubated with primary antibodies in 5% BSA at 4°C overnight. After washing three times with 1×PBS, primary antibodies were revealed using Alexa Fluor 488 donkey anti-rabbit antibody or Alexa Fluor 594 donkey anti-goat antibody (1:400) in 5% BSA for 30 minutes at 37°C. Images were captured using a confocal laser scanning microscope (LSM 780, Zeiss, Germany).

### 3.1. Western blot

Total proteins were extracted from HUVECs using RIPA buffer supplemented with protease inhibitors. Proteins were separated by 10% SDS-PAGE (Sigma-Aldrich, USA) and transferred to nitrocellulose membranes (GE Healthcare Life sciences, USA). Membranes were blocked with 5% non-fat milk at room temperature (RT) for 1 hour, then incubated with primary antibodies at 4°C overnight, followed by HRP-conjugated secondary antibodies (diluted at 1: 5000) at RT for 2 hours. The blots were detected by ECL chemiluminescence substrate (Thermo Scientific, Rochford, IL, USA).

### 3.2. LPS-Induced Endothelial Permeability Model Construction and Rescue Experiments

Cell Culture Model: HUVECs were treated with LPS (1 mg/mL) for 8 hours to induce endothelial permeability. For rescue experiments, cells were pretreated with NAV2729 (10 μM), an inhibitor of ARF6 , for 1 hour before LPS exposure.

Mouse Model: Mice were challenged with LPS ( E. coli O111:B4, 10 mg/kg, intraperitoneally) for 6 hours to induce endothelial permeability. For rescue experiments, mice received TJ-M2010-5 (50 mg/kg), an inhibitor of MyD88, daily for 7 days prior to LPS challenge.

### 3.3. GST Pulldown Assay for ARF6 Activation

HUVECs transfected with Scramble or G3BP1 siRNA were deprived of serum for 16 hours and then exposed to PBS or LPS (1 µg/ml) for the specified durations. The cells were lysed using buffer E (50 mM Tris–HCl, 1% Nonidet P-40, 100 mM NaCl, 10% glycerol, 5 mM MgCl2, and protease inhibitors, pH 7.4). The lysates were centrifuged for 10 minutes at 10,000 g at 4 ° C. GST-GGA3 fusion proteins bound to glutathione-Sepharose 4B beads were added to the supernatant and rotated at 4°C for 1 hour. After washing, the proteins were eluted in SDS-sample buffer with 5% β-mercaptoethanol by heating at 65°C for 15 minutes. The samples were separated on a 12% SDS-PAGE gel, and a Western blot analysis was performed.

### 3.4. Reverse transcription PCR (RT-PCR) and quantitative real-time RT-PCR (qRT-PCR)

Total RNA was isolated using FreeZol Reagent (Vazyme Biotech Co. Ltd, Cat no. R701-01, China), and cDNA was synthesized using HiFiScript cDNA Synthesis Kit (Cwbiotech, cat no. CW2569M, China) according to the instructions. RT-PCR was conducted using TaKaRa Ex Taq (RR001A, TaKaRa). qRT-PCR was performed using a TB Green™ Premix Ex Taq Kit (RR420A, TaKaRa) on an ABI QuantStudio DX real-time PCR machine (ABI, USA). We evaluated mRNA expression levels using the comparative threshold cycle method (2 −ΔΔ Ct) with GAPDH as an internal reference. The list of primers is provided in Additional file 1: Table S1.

### 3.5. RNA coimmunoprecipitation

RNA coimmunoprecipitation was conducted using an EZ-Magna RIP Kit (17–701, Millipore) following the manufacturer’s instructions. Briefly, HUVECs in the logarithmic growth phase were collected and suspended in RIP Lysis Buffer. After incubating for 30 minutes, the samples were centrifuged at 2500 g for 10 minutes at 4° C. Magnetic beads bound to anti-G3BP1 antibody or IgG for pre-clearance were rinsed with RIP Wash Buffer. The mixture was incubated overnight at 4°C with rotation. After isolating the magnetic beads, the supernatant was discarded. Then, 500 µl of RIP Wash Buffer was added to cleanse the beads. The mixture was vigorously mixed and the resulting precipitate was collected. A volume of 10 µl of the cell lysate supernatant was designated as the "Input" and stored temporarily at a temperature of -80°C. Subsequently, 150 µl of Proteinase K Buffer was added to the precipitate. A combination of RIP Wash Buffer, 10% SDS, and Proteinase K was added to the “Input” sample after freezing. This was followed by a 30 minute incubation at 55° C with shaking to facilitate protein digestion. RNA extraction was performed using the RNA-easy Isolation Reagent to obtain total RNA for qPCR analysis. The primer sequences for qPCR can be found in Additional file 1: Table S1.

### 3.6. RNA stability assay

HUVECs were transfected with Scramble or G3BP1 siRNA for 48 hours, followed by treatment with 5μg/mL actinomycin D (Sigma, St. Louis MO, USA) for 2, 4, or 6 hours. Total RNA was extracted using FreeZol Reagent, and mRNA levels were assessed by real-time PCR with GAPDH as an internal control. The mRNA half-life was calculated from the decay rate. Primers are in Additional file 1: Table S1.

### 3.7. Statistical analysis

Data are presented as mean ± standard deviation. All statistical analyses were performed using GraphPad Prism software. Data were first assessed for normality using the Shapiro-Wilk test and for equal variance using the Brown-Forsythe test. Comparisons between groups were made using an unpaired Student’s t-test or a two-way ANOVA, as appropriate. Statistical significance was set at P < 0.05. Significant differences between groups are indicated as follows: * P < 0.05, ** P < 0.01, *** P < 0.001.

## 3. Results

### G3BP1 maintains vascular endothelial permeability, particularly under LPS-induced inflammatory conditions

To determine the role of *G3BP1* in vascular endothelial integrity, we assessed endothelial permeability in endothelial-specific *G3bp1* knockout mice and *G3BP1*-depleted HUVECs.

We generated endothelium-specific *G3bp1* knockout mice (*G3bp1 ^flox/flox^; Cdh5*: *Cre, G3bp1* cKO; Supplementary Fig. 1) and assessed endothelial permeability in these mice under both resting and LPS-induced inflammatory conditions using the Miles assay, lung water analysis, and histological evaluation.

Under resting conditions, we observed no significant differences in vascular permeability, as measured by Evans blue dye (EBD) leakage and lung wet/dry weight ratio, between *G3bp1* cKO and control mice (Fig. 1A-1E). However, histological analysis revealed subtle localized changes within the lung tissue of *G3bp1* cKO mice, including increased inflammatory cell infiltration and thicker, more irregular alveolar walls compared to control mice (Fig. 1F).

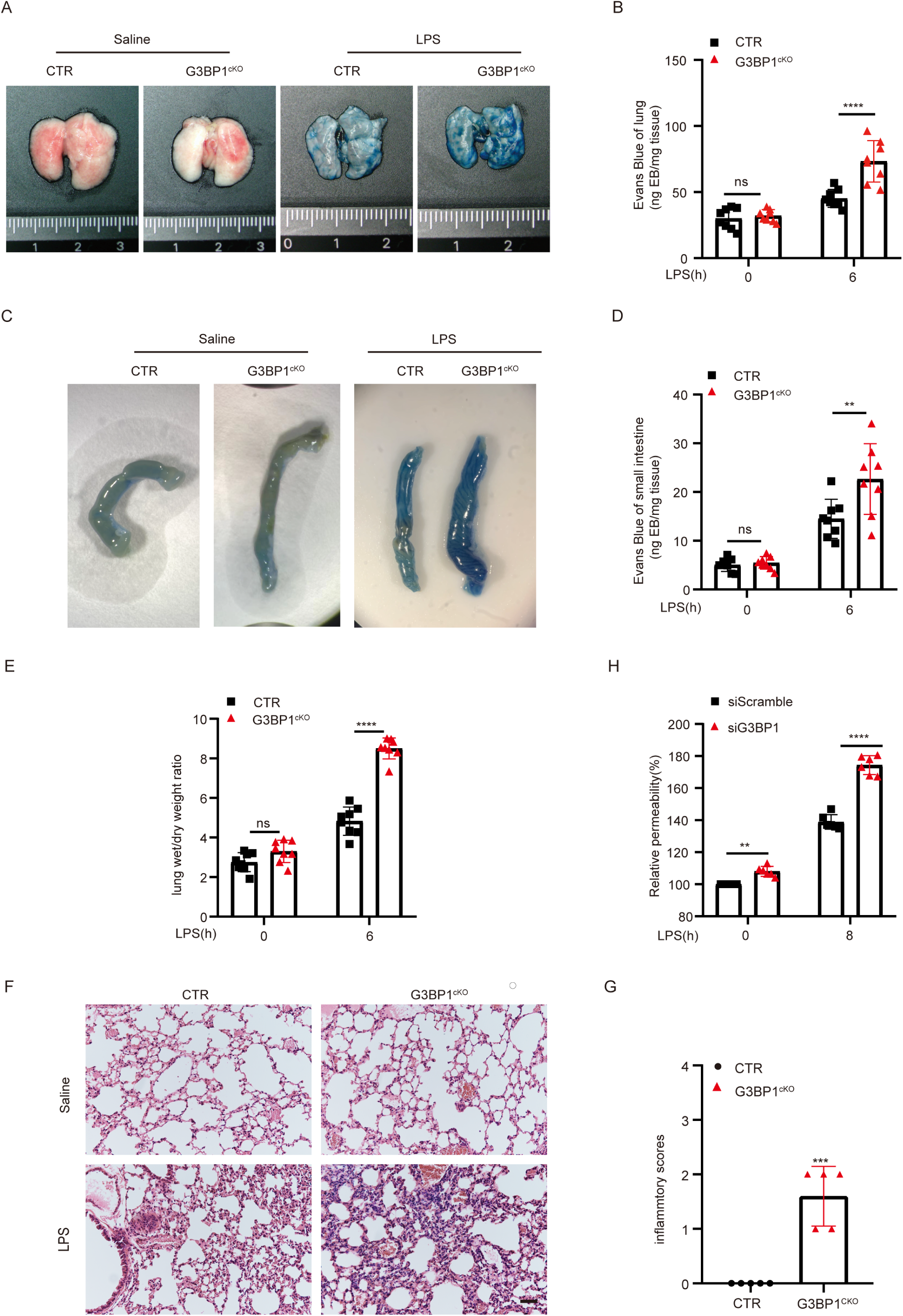

In contrast, under LPS-induced inflammatory conditions, *G3bp1* cKO mice exhibited significantly increased vascular permeability. EBD leakage was approximately 62% higher in the lung and 56% higher in the small intestine of *G3bp1* cKO mice than in control mice (Fig. 1B, n=8, P < 0.05; Fig. 1D, n=8, P < 0.05). The lung wet/dry weight ratio was also significantly elevated in *G3bp1* cKO mice compared to control mice (8.50±0.53 vs. 4.83±0.71, n=8 per group, P < 0.05) (Fig. 1E). Consistent with these findings, *G3bp1* cKO mice showed more pronounced PMN sequestration and alveolar wall thickening than control mice (Fig. 1F and 1G).

To confirm the role of G3BP1 in endothelial cell permeability, we knocked down *G3BP1* in HUVECs using siRNA, then examined cell permeability using *in vitro* FITC-dextran permeability assay. While G3BP1 knockdown led to a modest increase in permeability under resting conditions, LPS stimulation significantly amplified this effect, resulting in a 26% increase in permeability compared to that in control cells [Fig. 1H].

In summary, these results indicate that loss of G3BP1 increased endothelial permeability, particularly under LPS-induced inflammatory condition, both in vivo and in vitro.

### Loss of G3BP1 impairs Vascular Endothelial Barrier Integrity

To investigate whether the increase in vascular permeability was associated with impaired endothelial barrier integrity, we examined AJ and TJ in lung vascular endothelial cells of *G3bp1* cKO mice using transmission electron microscopy (TEM) and immunostaining, as well as in *G3BP1*-depleted HUVECs using immunostaining.

First, we directly observed endothelial junction integrity in lung vascular endothelial cells using TEM. Under resting conditions, control mice exhibited continuous, electron-dense junctional complexes between adjacent pulmonary endothelial cells, indicative of intact barrier integrity (Fig. 2A, red arrows). In contrast, *G3bp1* cKO mice displayed a disruption of endothelial junctions, characterized by intercellular gaps and less electron-dense junctional complexes (Fig. 2A, red arrows). Following LPS administration, control mice showed widening of cell junctions (Fig. 2A, red arrows). However, *G3bp1* cKO mice exhibited a dramatic exacerbation of junctional disruption. Quantitative analysis of TEM images revealed a significant decrease in the percentage of pulmonary vessels with intact endothelial junctions in *G3bp1* cKO mice compared to control mice, both under basal conditions and after LPS challenge (Fig. 2B).

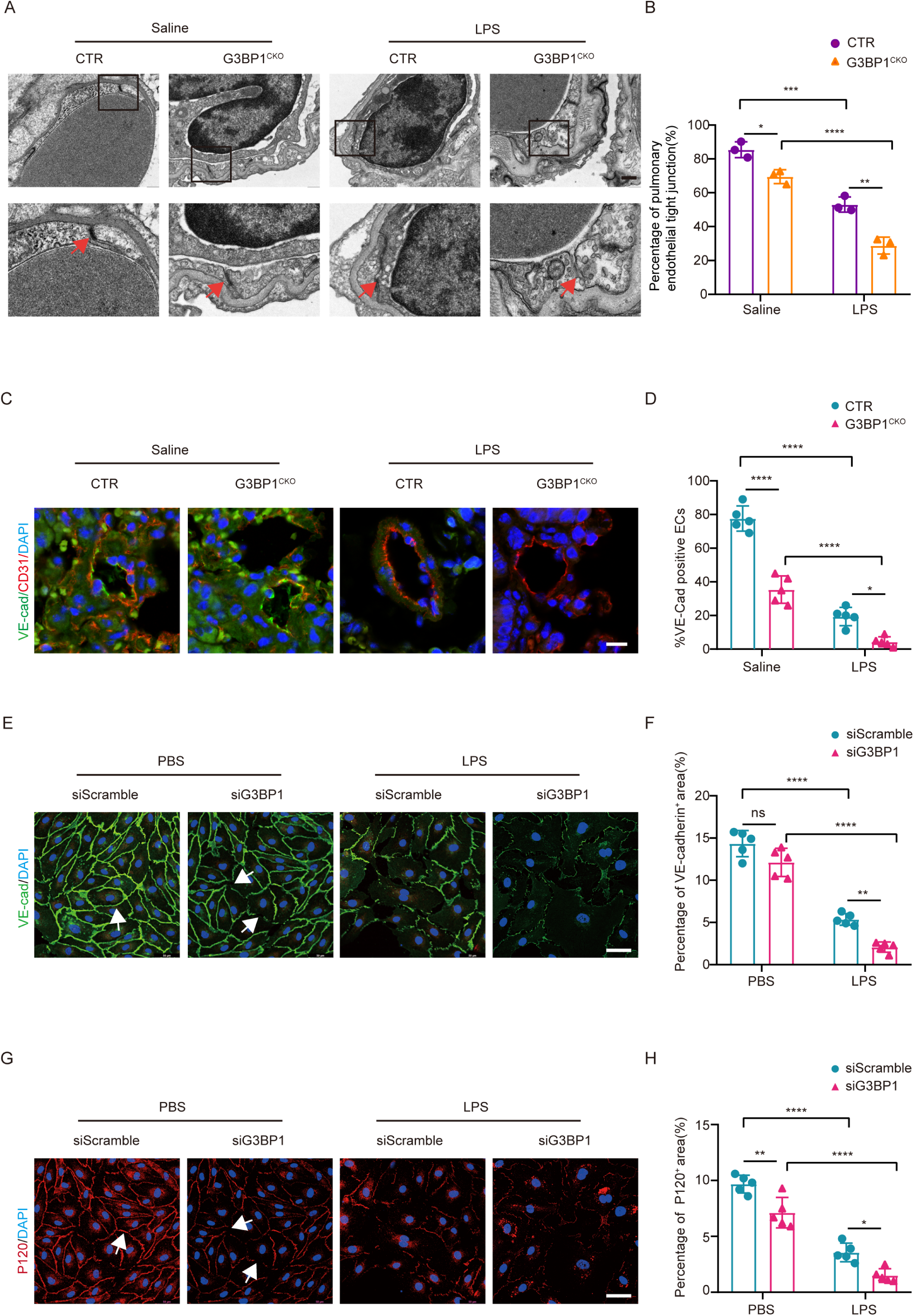

Next, we examined the localization of the VE-cadherin and p120 using immunostaining. Under resting conditions, HUVECs exhibited a continuous, well-defined VE-cadherin staining pattern along cell-cell borders, indicating intact adherens junctions (Fig. 2E and 2G). In contrast, G3BP1 depletion in HUVECs resulted in a reduction in continuous VE-cadherin and p120 staining at cell-cell borders, even in the absence of LPS stimulation (Fig. 2E and 2G). consistent with results in HUVECs, *G3bp1* cKO mice exhibited less endothelial cells with VE-cadherin staining than control mice (Fig. 2C and 2D). Under LPS-induced inflammatory conditions, HUVECs exhibited disrupted VE-cadherin organization. This disruption manifested as a more irregular and discontinuous staining pattern with gaps and reduced intensity at cell-cell borders (Fig. 2C and 2D). *G3BP1* depletion significantly exacerbated this disruption. *G3bp1* cKO mice and G3BP1-depleted HUVECs displayed a near-complete loss of VE-cadherin staining at cell-cell borders following LPS challenge (Fig. 2C and 2D).

In summary, these findings indicate that G3BP1 deficiency impairs vascular endothelial barrier integrity, particularly under inflammatory conditions.

### G3BP1 Regulates VE-Cadherin and p120 Expression in Vascular Endothelial Cells

Given the observed impairment of endothelial barrier integrity in our previous experiments, we asked whether G3BP1 regulates the expression of AJ and TJ proteins. We examined the protein and mRNA levels of key AJ and TJ components under conditions of *G3BP1* depletion and overexpression using western blot and qRT-PCR analyses, respectively.

We observed that *G3BP1* depletion significantly reduced the protein expression of AJ components, including VE-cadherin and p120, under both resting and LPS-challenged conditions in mouse lung tissues (Fig. 3A and 3B) and HUVECs (Fig. 3C and 3D). Similarly, *G3BP1* depletion led to a significant decrease in the protein levels of TJ proteins, specifically ZO-1 and Claudin-5, in HUVECs under both conditions (Supplementary Fig. 2A). Notably, in contrast to the reductions observed in protein levels, the mRNA levels of p120 and VE-cadherin were decreased in *G3BP1*-depleted HUVECs (Fig. 3E), while the mRNA levels of ZO-1 and Claudin-5 remained unchanged (Supplementary Fig. 2B). These findings suggest that G3BP1 regulates the expression of AJ and TJ proteins through distinct mechanisms.

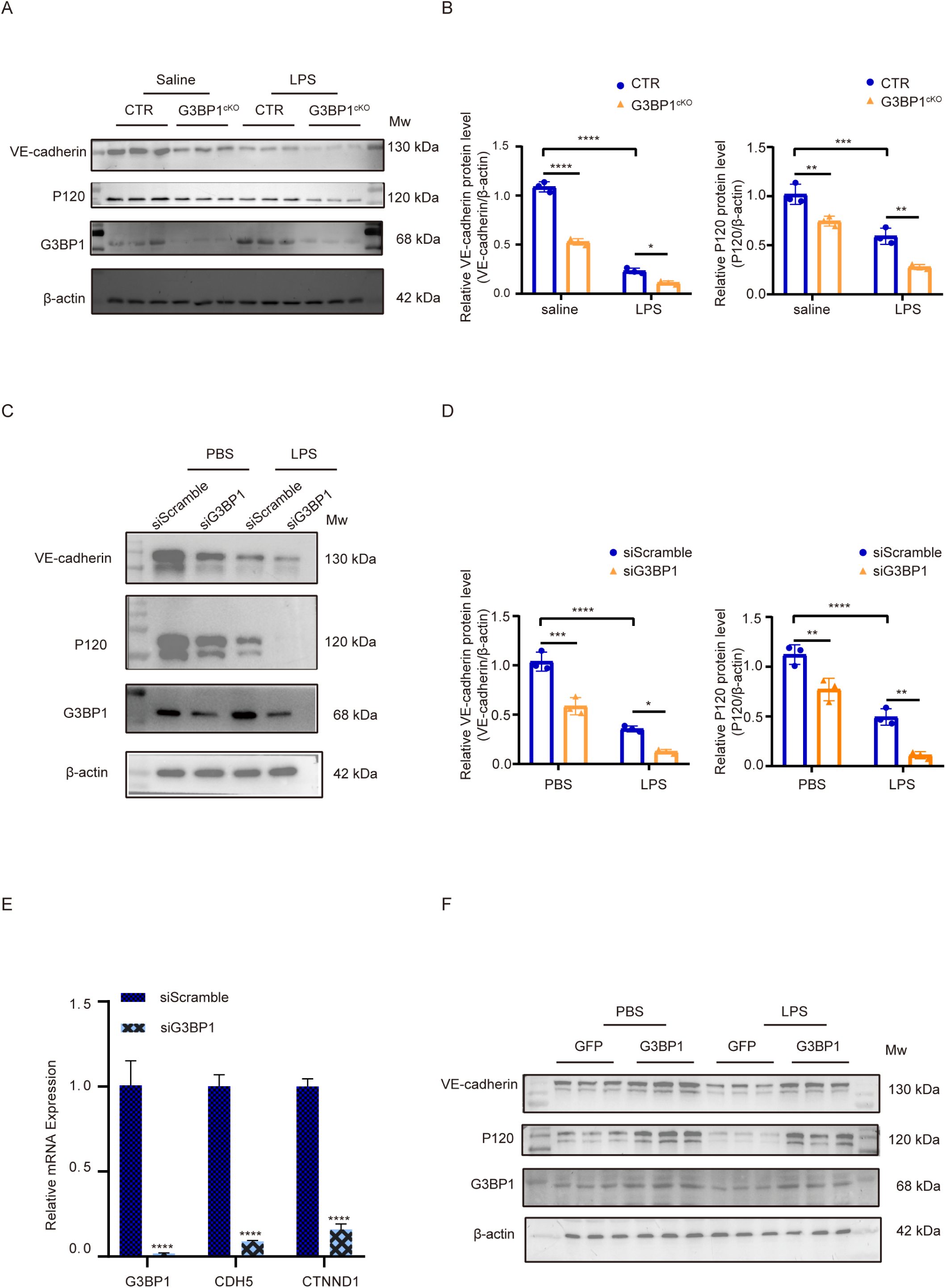

To better understand the specific effects of G3BP1 on junctional protein expression, we overexpressed *G3BP1* in HUVECs. Under resting conditions, *G3BP1* overexpression evidently increased the protein levels of AJ components VE-cadherin and p120 (Fig. 3F), while did not alter protein levels of TJ including ZO-1 and Claudin-5 (Supplementary Fig. 2C). Under LPS-induced inflammatory conditions, LPS challenge reduced protein levels of both AJ and TJ (Fig. 2F). Notably, *G3BP1* overexpression restored VE-cadherin and p120 expression but did not prevent the LPS-induced decrease in ZO-1 and Claudin-5 (Fig. 3F and Supplementary Fig. 3C).

In summary, our results indicate that G3BP1 regulates the expression of AJ protein VE-Cadherin and p120 in vascular endothelial cells.

### Loss of G3BP1 activates MYD88-ARNO-ARF6 Signaling under LPS-induced inflammatory conditions

To investigate the mechanism by which Loss of G3BP1 increases endothelial permeability under LPS-induced inflammatory conditions, we focused on the MYD88-ARNO-ARF6 signaling pathway. This pathway, activated by LPS, is known to trigger VE-cadherin endocytosis and degradation, leading to the disruption of endothelial barrier integrity[14]. We investigated whether *G3BP1* depletion, under LPS-induced inflammatory conditions, enhanced the activation of the MYD88-ARNO-ARF6 signaling pathway.

First, we examined the effects of *G3BP1* depletion on the expression of key components within this pathway. Under resting conditions, *G3BP1* depletion significantly increased the protein levels of MyD88, ARNO, and ARF6 in HUVECs (Fig. 4A). Under LPS-induced inflammatory conditions, we observed the similar increase in both HUVECs (Fig. 4A) and mouse lung tissues (Fig. 4F-4I).

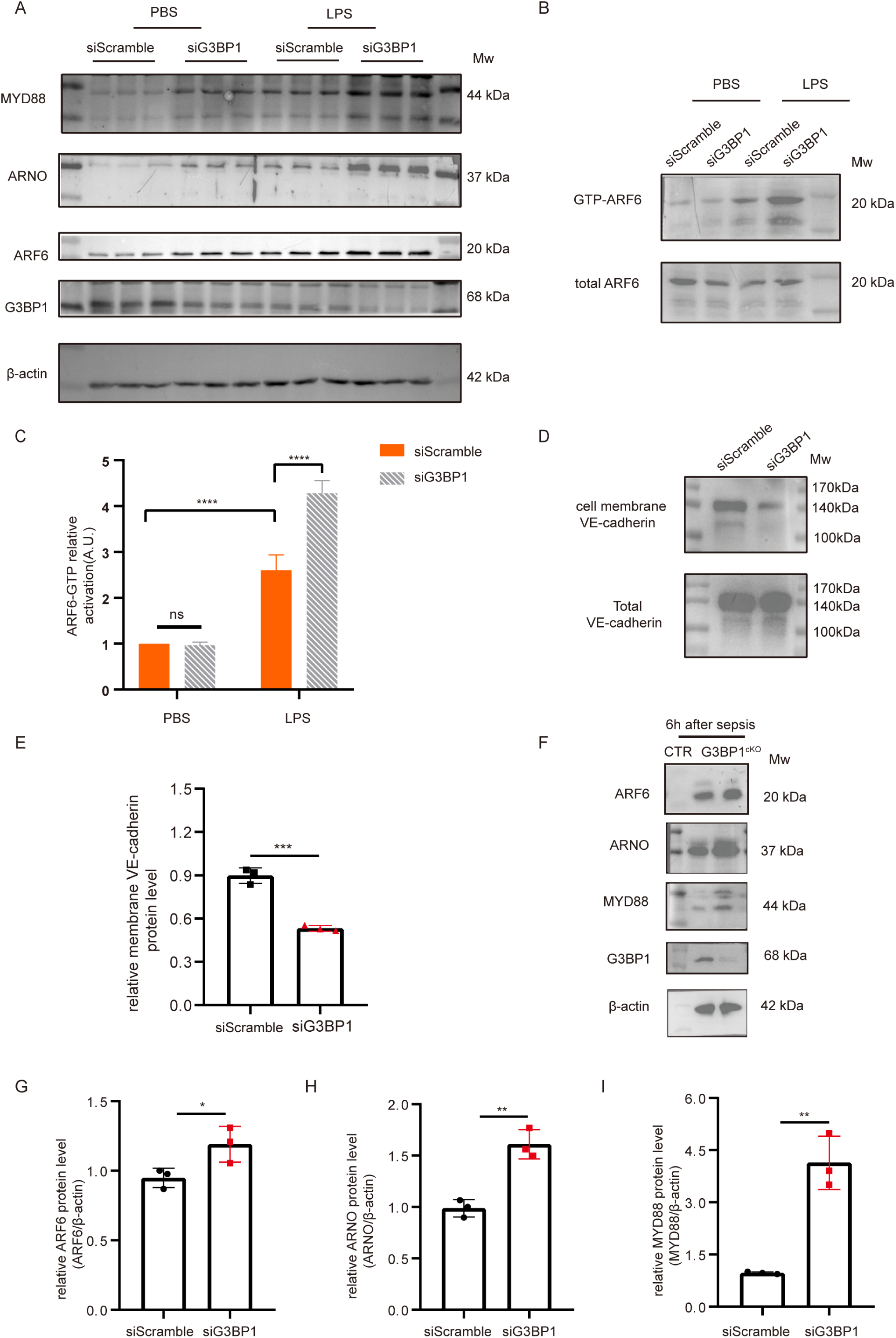

Next, we assessed the activation status of ARF6 by measuring ARF6-GTP levels. Under resting conditions, no significant difference in ARF6 activation was detected between *G3BP1*-depleted and control cells (Fig. 4B). LPS challenge significantly increased ARF6-GTP levels in control HUVECs, and *G3BP1*-depleted HUVECs exhibited higher ARF6-GTP levels than control HUVECs (Fig. 4B and 4C).

Finally, we examined the effect of *G3BP1* knockdown on membrane-bound VE-cadherin, which is associated with ARF6-mediated internalization. *G3BP1* knockdown decreased membrane VE-cadherin levels (Fig. 4D and 4E, n = 3, P < 0.05) under LPS-induced inflammatory conditions, suggesting increased internalization.

In summary, our findings indicate that *G3BP1* depletion enhanced the activation of the MYD88-ARNO-ARF6 signaling pathway under LPS-induced inflammatory conditions.

### G3BP1 Deficiency Enhances Endothelial Permeability via the MYD88-ARNO-ARF6 Pathway

To determine whether the activated MYD88-ARNO-ARF6 signaling pathway contributes to the increased endothelial permeability in *G3BP1*-deficient cells and mice, we investigated the effects of pathway inhibition on barrier integrity and endothelial permeability.

In *G3bp1* cKO mice, inhibition of MyD88 using the chemical inhibitor TJM significantly reduced EBD leakage (Fig. 5A-5D) and restored the expression of VE-Cadherin and p120 protein in the lung under LPS-induced inflammatory conditions (Fig. 5G-5H, 5K-5L; 5O), compared to vehicle-treated *G3bp1* cKO mice (Fig. 5E-5F, 5I-5J; 5O). Similarly, in *G3BP1-*depleted HUVEC cells, *MYD88* knockdown decreased cell permeability (Fig. 6A), restored barrier integrity (Fig. 6C-6F), and protein levels of VE-Cadherin and p120 under LPS-induced inflammatory conditions (Fig. 6B), compared to control siRNA-treated G3BP1 knockdown cells. Consistent with MyD88 inhibition, knockdown of *ARF6* and inhibition of ARNO by NAV also significantly improved barrier integrity and endothelial permeability in *G3BP1*-deficient cells under LPS-induced inflammatory conditions (Fig. 6I-6J, 6K-6L).

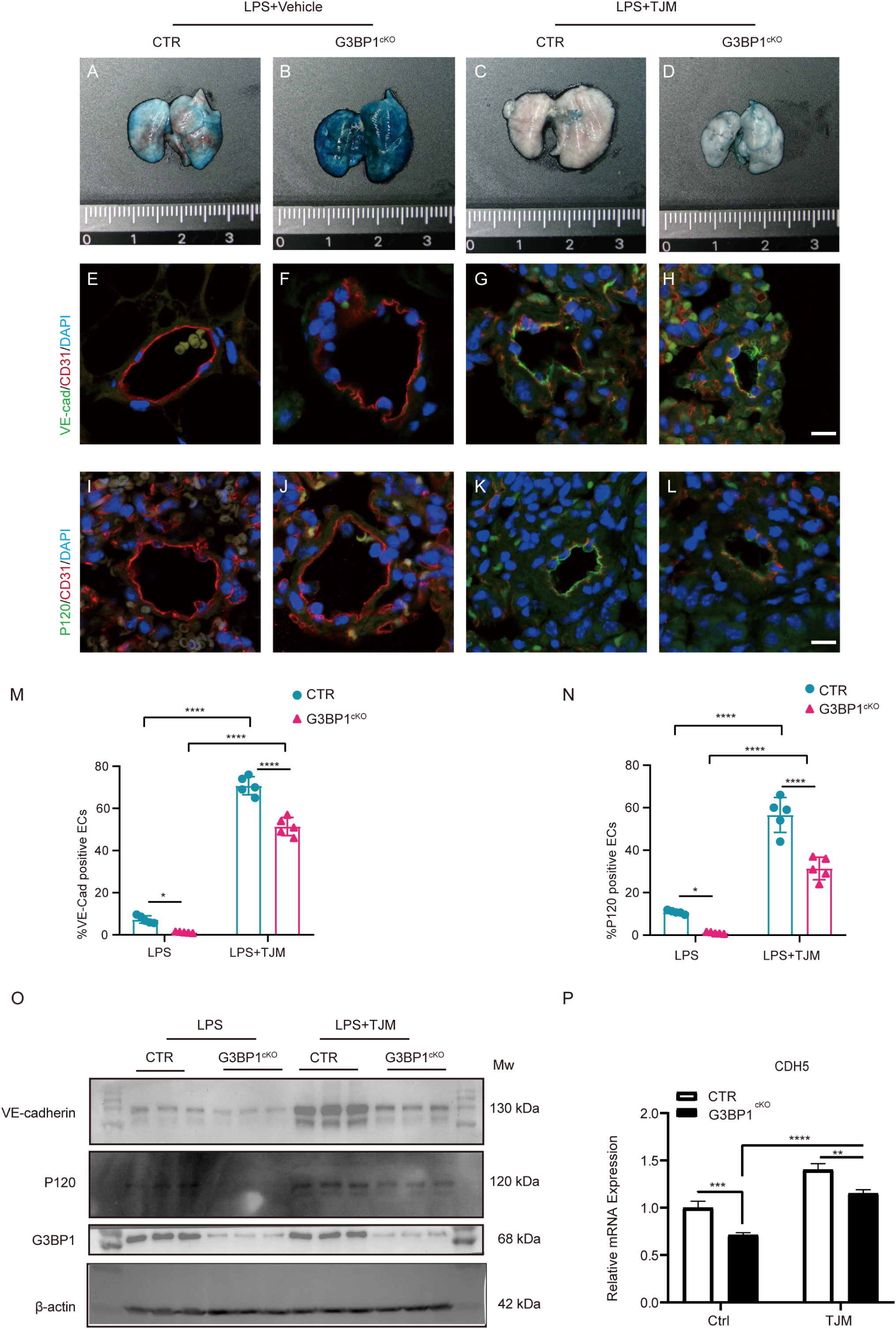

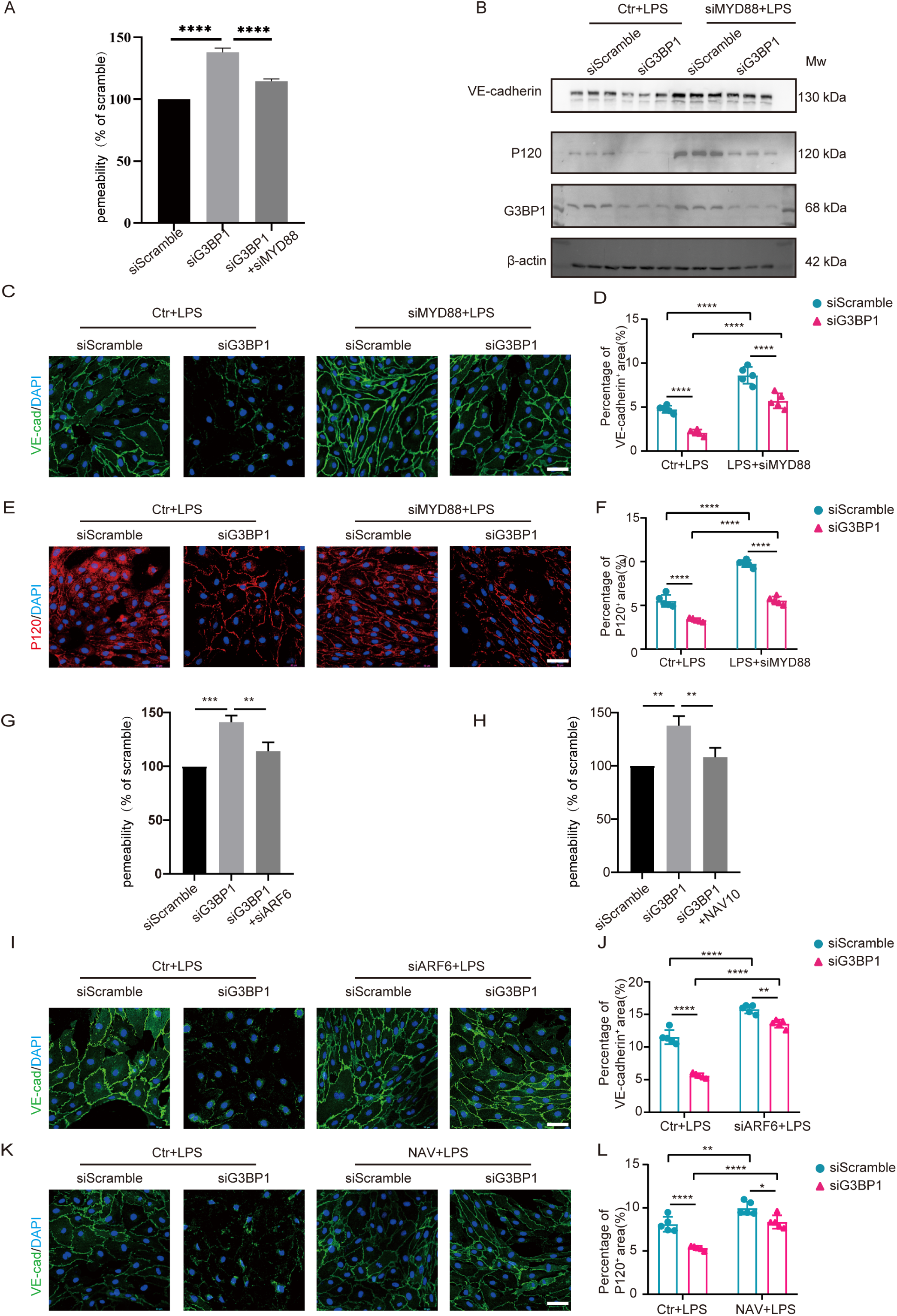

In summary, our results demonstrate that inhibiting the MYD88-ARNO-ARF6 signaling pathway effectively restores endothelial barrier integrity and reduces permeability in *G3BP1*-deficient cells and mice under LPS-induced inflammatory conditions.

### G3BP1 regulates MyD88 mRNA stability

G3BP1 is an RNA-binding protein known to bind mRNA and regulate its stability. To explore whether G3BP1 regulates the MYD88-ARNO-ARF6 signaling pathway post-transcriptionally, firstly, we assessed the mRNA expression of MYD88, ARNO, and ARF6 in *G3BP1* knockdown cells. Consistent with our protein-level findings, the mRNA levels of MYD88, ARNO, and ARF6 were obviously increased in *G3BP1* knockdown cells compared to controls (Fig. 7A, n =3, P < 0.05). We observed similar increases in mRNA levels in the lung tissues of *G3bp1* cKO mice compared to control mice (Fig. 6B, n =3, P < 0.05).

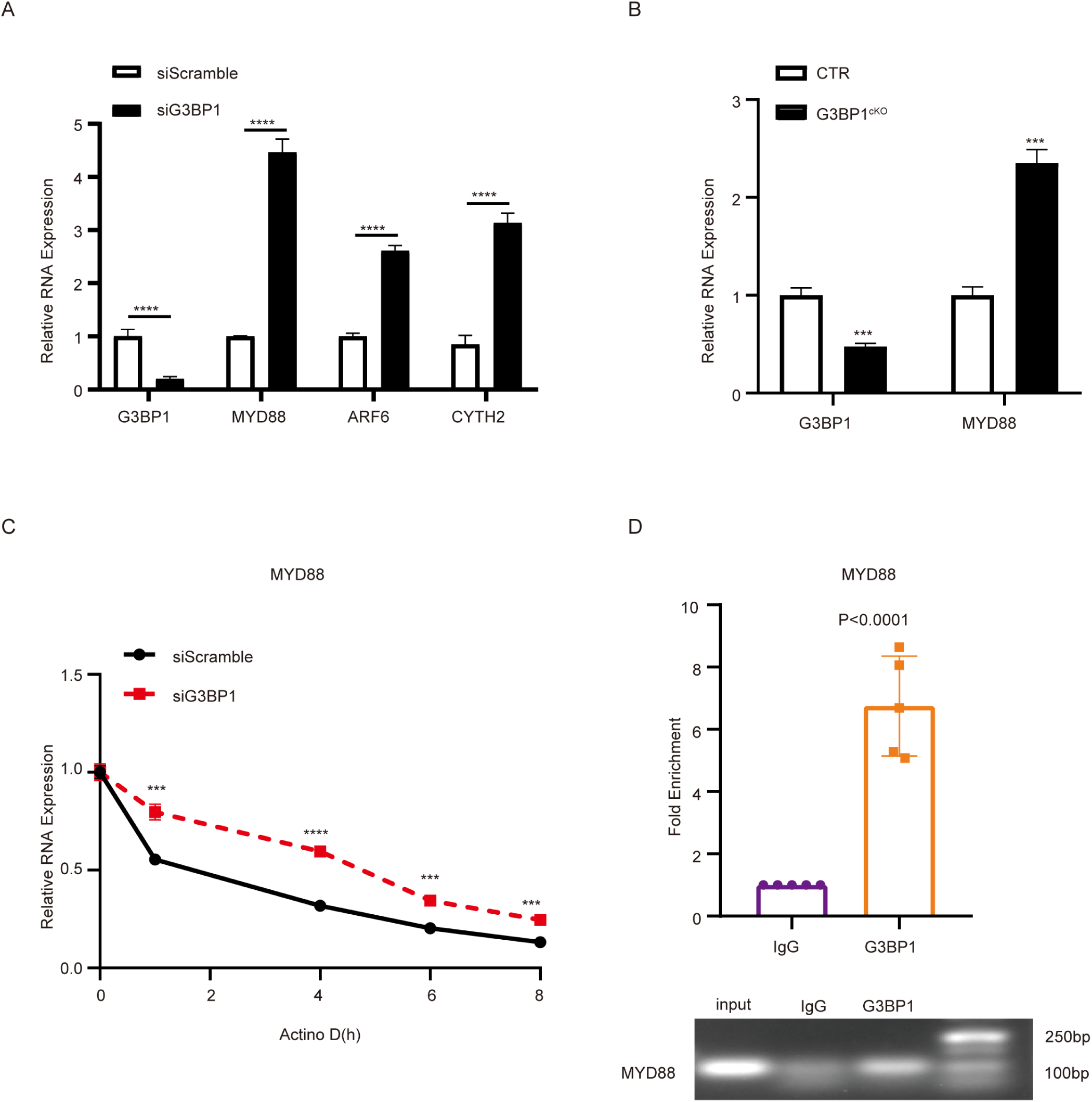

Next, we examined whether G3BP1 affected the stability of these mRNAs. In the presence of Actinomycin D, which inhibits transcription, *MYD88* mRNA exhibited a slower decay in *G3BP1* knockdown cells than in control cells, indicating increased mRNA stability (Fig. 7C, n = ?, P < 0.05). However, no significant differences in mRNA stability were detected for *ARNO* and *ARF6* between *G3BP1* knockdown and control cells (Supplementary fig. 3A-3B, n = 2, P > 0.05).

To ask whether G3BP1 can bind to *MYD88* mRNA, we conducted RIP-PCR using a G3BP1-specific antibody. We observed a 6.2-fold enrichment of *MYD88* mRNA compared to the IgG control (Fig. 7D, n = 5, P < 0.05). In contrast, we found that no significant enrichment for *ARNO* and *ARF6* mRNA between G3BP1 and IgG antibody (Supplementary fig. 3C-3D). As a positive control, *CTNNB1* mRNA, a known G3BP1 target, showed significant enrichment, whereas *GAPDH* mRNA, used as a negative control, did not show enrichment (data not shown).

We also explored the possibility that G3BP1 directly regulates mRNA stability of AJ including VE-Cadherin and p120. We examined the stability of their corresponding mRNAs (*CDH5* and *CTNND1*, respectively). Actinomycin D experiments revealed a significantly faster decay rate for both *CDH5* and *CTNND1* transcripts in *G3BP1* knockdown cells compared to control cells (Fig. 8E-F), indicating a reduction in mRNA stability (Fig. 8E and 8F).

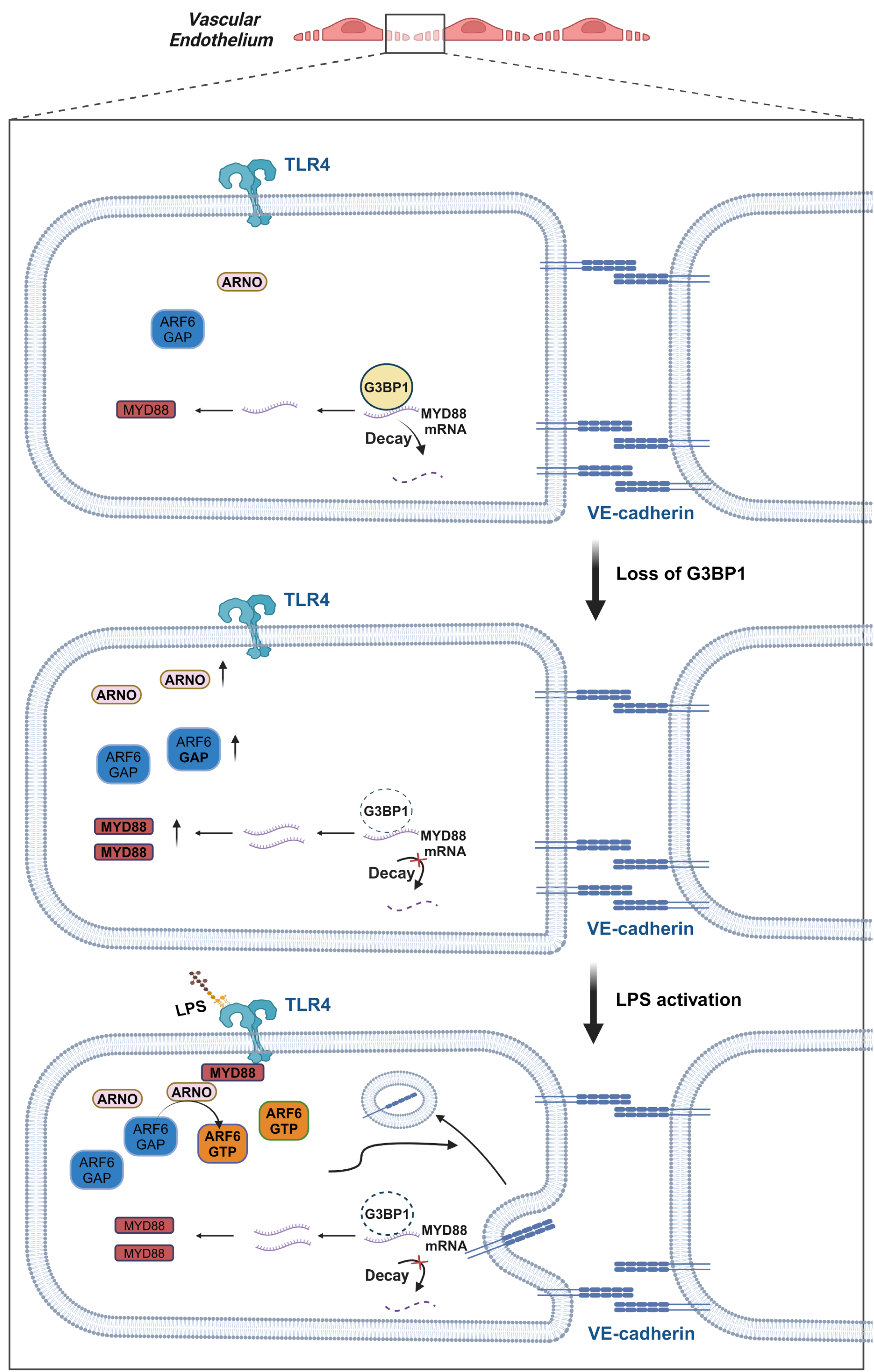

We further examined whether G3BP1 can directly bind to *CDH5* and *CTNND1* mRNA using RIP-qPCR. G3BP1 can bind to the mRNA of *CDH5* but not *CTNND1* (Supplementary Fig. 3G and 3H), suggesting that G3BP1 directly regulates *CDH5* mRNA stability.

Taken together, these results suggest that G3BP1 binds to *MYD88* mRNA and regulates its stability, thereby regulating the MYD88-ARNO-ARF6 signaling pathway.

## 4. Discussion

This study shows that G3BP1 maintains vascular endothelial barrier integrity by negatively regulating the MYD88-ARNO-ARF6 signaling pathway. In our experiments, loss of G3BP1 in vascular endothelial cells of both mice and humans decreases the expression of AJs and TJs, compromises barrier integrity, and increases permeability, especially under LPS-induced inflammatory conditions (Fig. 1, Fig. 2 and Fig. 3). These effects are mediated, at least in part, through G3BP1’s ability to bind to MYD88 mRNA and regulate its stability (Fig. 7). Reducing the increased expression of MYD88 or ARF6, or inhibiting ARNO activity, restores the expression of VE-Cadherin and p120, barrier integrity, and reduces permeability in *G3BP1*-deficient cells and mice (Fig. 5 and Fig. 6). Our evidence identify G3BP1 as a novel molecular player in endothelial barrier regulation (Fig. 8).

The MYD88-ARNO-ARF6 signaling cascade plays a crucial role in vascular stability and its activation by IL-1 β or LPS increase vascular endothelial permeability[13, 14]. Our findings build upon this knowledge. Firstly, we showed that loss of G3BP1 increased the mRNA and protein levels of MYD88, ARNO, and ARF6 under resting conditions (Fig.4A, 4F and 7A). Upon LPS challenge, ARF6 activity was significantly enhanced, resulting in MYD88-ARNO-ARF6 signaling activation and endothelial permeability increase (Fig.4B and 4C). Moreover, Inhibiting the activation of MYD88-ARNO-ARF6 signaling by reducing the levels of MYD88 or ARF6, or inhibiting ARNO activity, reverses the impairment of endothelial barrier integrity and permeability increase caused by loss of *G3bp1* (Fig. 5 and Fig. 6). Secondly, transient activation of MYD88-ARNO-ARF6 signaling increases endocytosis and internalization of VE-cadherin[13], while prolonged activation decreases both mRNA and protein levels of VE-cadherin[13, 33]. Correcting the activation of MYD88-ARNO-ARF6 signaling can restore the protein expression of VE-cadherin and p120 [34]. We showed that inhibiting MYD88 activity or reducing MYD88 levels can restore the expression of VE-cadherin and p120 caused by loss of *G3bp1* (Fig. 5O, Fig. 6B). Thirdly, mechanistically, G3BP1 promoted *MYD88* mRNA decay through binding to its mRNA (Fig. 7). Loss of *G3bp1* disrupts this regulation, as a result, more *MYD88* mRNA and protein accumulate, which underlie over-activation of MYD88-ARNO-ARF6 signaling by LPS (Fig. 8). Our results are consistent with the established role of MYD88-ARNO-ARF6 signaling in endothelial barrier function and G3BP1’s regulatory role as an RNA-binding protein.

In addition to its effects on the MYD88-ARNO-ARF6 pathway, we also explored the possibility that G3BP1 directly regulates the expression of AJ and TJ proteins. Our results showed that G3BP1 bound to the *CDH5* mRNA and decreased its stability (Supplementary Fig. 3E and 3F). This finding suggests that G3BP1 regulates endothelial barrier integrity through multiple mechanisms.

We observed a discrepancy between membrane VE-cadherin levels and ARF6 activation state in *G3BP1* knockdown HUVEC cells under resting conditions. Membrane VE-cadherin was significantly decreased, but ARF6 activation was not altered. The decrease of membrane VE-cadherin cannot be solely attributed to the decreased total protein levels of VE-cadherin, as the membrane and total VE-cadherin levels decreased disproportionately. *G3BP1* depletion may affect other mechanisms involved in VE-cadherin trafficking or stabilization at the membrane, independent of ARF6 activation.

Furthermore, the simultaneous increase in the mRNA levels of *MYD88*, *ARNO*, and *ARF6* upon loss of *G3bp1* raises intriguing questions. While we found that G3BP1 directly binds to *MYD88* mRNA and regulates its stability, it does not appear to directly interact with *ARNO* or *ARF6* mRNA (Supplementary Fig. 3A-3D). Therefore, we speculate that G3BP1 may indirectly affect the expression of ARNO and ARF6, possibly by modulating other RNA-binding proteins or microRNAs that target these genes.

Our study adds to the growing body of evidence highlighting the importance of post-transcriptional regulation by RBPs in endothelial barrier function. Although numerous RBPs have been identified in vascular endothelial cells, only a few have been thoroughly investigated for their roles in regulating endothelial barrier function. Among them, two RBPs, QKI and TDP-43, have been shown to regulate AJ components. QKI binds to the 3’UTRs of VE-cadherin and β-catenin mRNA, ensuring sufficient translation to restrict vascular permeability [15]. Similarly, TDP-43 promotes nuclear localization and signaling of β-catenin, a protein that interacts with VE-cadherin [35]. In addition to its similar role in directly regulating VE-cadherin mRNA, *CHD5,* like QKI, G3BP1 regulates the MYD88-ARNO-ARF6 pathway, primarily by controlling *MYD88* mRNA stability. This finding expands our understanding of the diverse mechanisms by which RBPs can influence endothelial barrier integrity.

Moreover, our study identifies G3BP1 as a novel negative regulator of the MYD88-ARNO-ARF6 pathway. This pathway is subject to complex regulation, with several known negative regulators, including IRAK-M, SOCS1, miR-146a, and CYLD, acting at different levels (e.g., kinase activity, receptor signaling, gene expression, and ubiquitination). G3BP1 adds a new layer of complexity to this regulation by acting at the level of mRNA stability. Further studies can be done to investigate whether G3BP1 interacts with or influences other negative regulators to fine-tune the activity of the MYD88-ARNO-ARF6 cascade.

The discovery of G3BP1 as a regulator of the MYD88-ARNO-ARF6 cascade has broad implications for understanding and potentially treating diseases associated with vascular instability and inflammation. Dysregulation of AJ dynamics and vascular permeability is a hallmark of various pathological conditions, such as inflammation, sepsis, and cancer metastasis. Our study suggests that targeting RNA-binding proteins or their downstream effectors may represent a promising therapeutic strategy for these diseases.

One limitation of our study is that we only explored G3BP1’s role in vascular integrity in endothelial cells. Future studies could explore G3BP1’s role in other cell types involved in vascular integrity, such as vascular smooth muscle cells. Furthermore, while our findings in cellular and mouse models strongly suggest a critical role for G3BP1 in maintaining endothelial barrier integrity, further investigation is warranted to determine its relevance to human vascular disease. Specifically, analyzing *G3BP1* expression levels in patient cohorts could provide valuable insights. For instance, correlating G3BP1 levels with disease severity markers (e.g., vascular permeability, inflammatory markers) in conditions like acute respiratory distress syndrome or sepsis could reveal whether G3BP1 serves as a potential prognostic indicator or therapeutic target in these settings.

In conclusion, our study demonstrates that G3BP1 maintains vascular endothelial barrier integrity by negatively regulating the MYD88-ARNO-ARF6 signaling pathway. These findings highlight the importance of post-transcriptional regulation by RNA-binding proteins in endothelial barrier function and suggest G3BP1 as a potential target for clinical intervention in permeability-associated diseases.

## Acknowledgment

This research was supported in part by the National Natural Science Foundation of China (82370509 and 82370510), Zhejiang Provincial Natural Science Foundation of China (LY24H020006), and Municipal-University Joint Funding Project for Special Topics in Basic and Applied Research (SL2022A03J01104)

## Abbreviation

AJs: adherens junction
TJs: tight junction
HUVECs: Primary Human Umbilical Vein Endothelial Cells

---

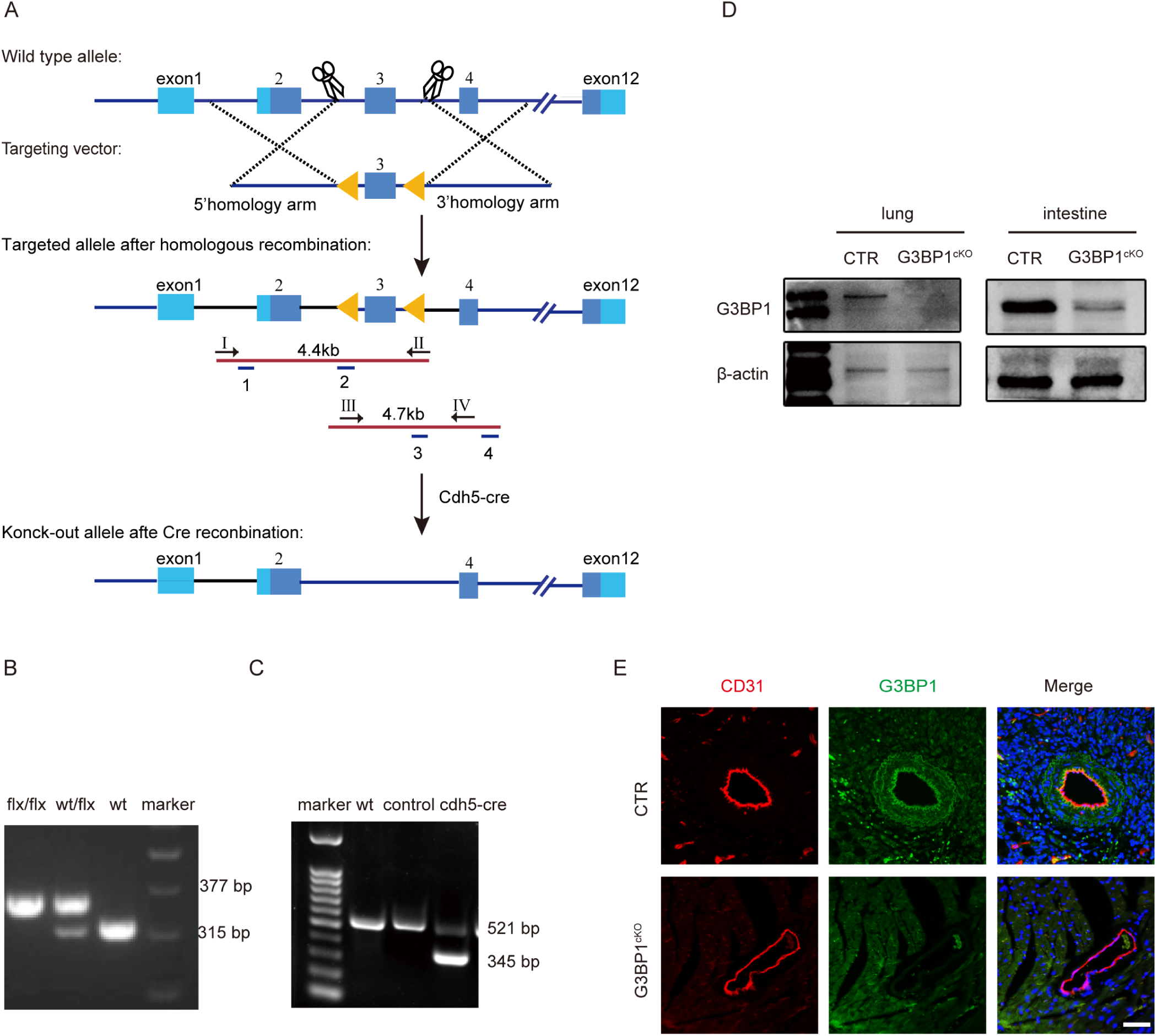

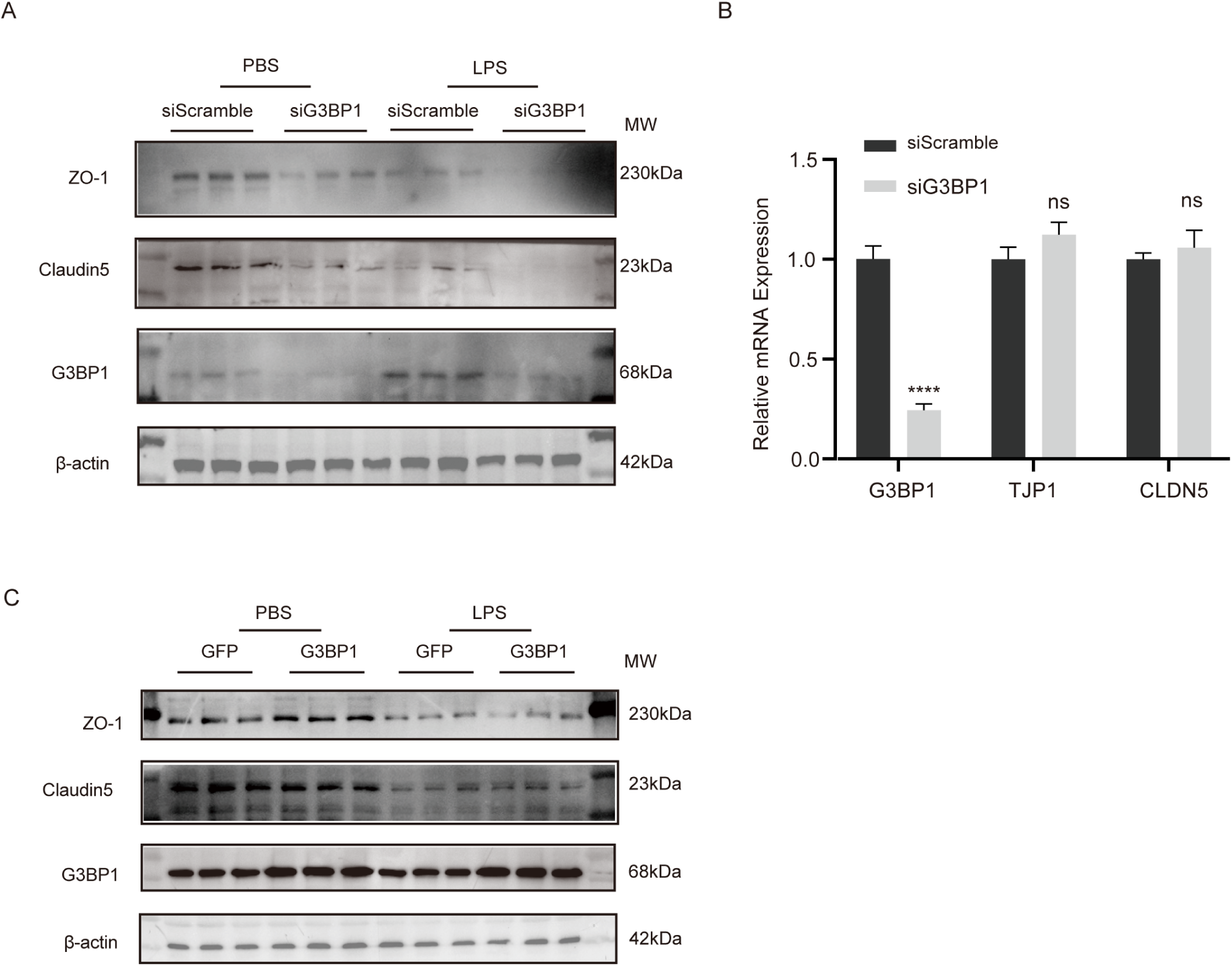

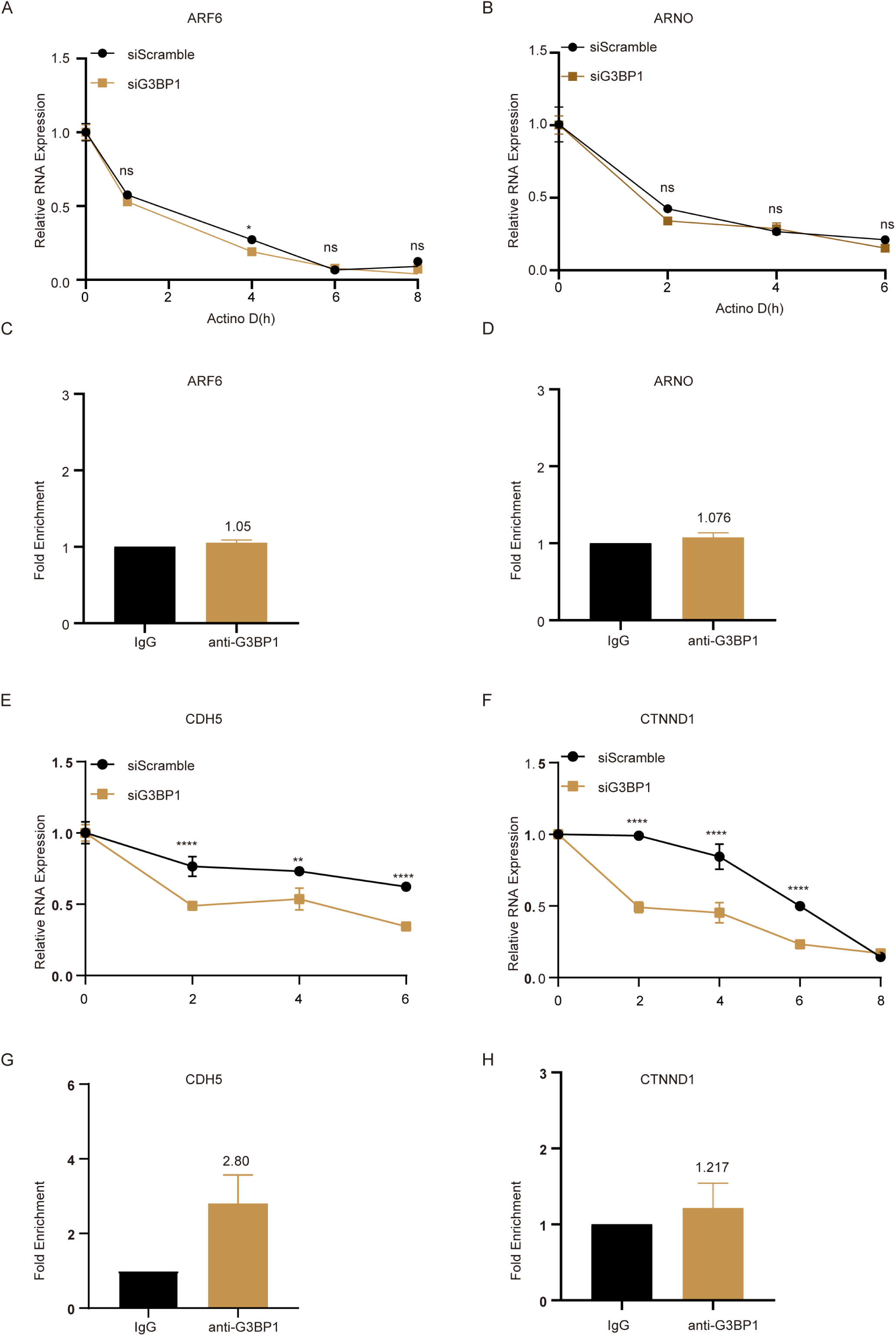

